# Effects of Mutations in the Receptor-Binding Domain of SARS-CoV-2 Spike on its Binding Affinity to ACE2 and Neutralizing Antibodies Revealed by Computational Analysis

**DOI:** 10.1101/2021.03.14.435322

**Authors:** Marine E. Bozdaganyan, Olga S. Sokolova, Konstantin V. Shaitan, Mikhail P. Kirpichnikov, Philipp S. Orekhov

## Abstract

SARS-CoV-2 causing coronavirus disease 2019 (COVID-19) is responsible for one of the most deleterious pandemics of our time. The interaction between the ACE2 receptors at the surface of human cells and the viral Spike (S) protein triggers the infection making the receptor-binding domain (RBD) of the SARS-CoV-2 S-protein a focal target for the neutralizing antibodies (Abs). Despite the recent progress in the development and deployment of vaccines, the emergence of novel variants of SARS-CoV-2 insensitive to Abs produced in response to the vaccine administration and/or monoclonal ones represents upcoming jeopardy. Here, we assessed the possible effects of single and multiple mutations in the RBD of SARS-CoV-2 S-protein on its binding energy to various antibodies and the human ACE2 receptor. The performed computational analysis indicates that while single amino acid replacements in RBD may only cause partial impairment of the Abs binding, moreover, limited to specific epitopes, some variants of SARS-CoV-2 (with as few as 8 mutations), which are already present in the population, may potentially result in a much broader antigenic escape. We also identified a number of point mutations, which, in contrast to the majority of replacements, reduce RBD affinity to various antibodies without affecting its binding to ACE2. Overall, the results provide guidelines for further experimental studies aiming at the identification of the high-risk RBD mutations allowing for an antigenic escape.

## Introduction

Recent start of vaccination campaigns in many countries allowed by the rapid development of several effective vaccines [1] gives hope for a forthcoming amelioration of the world pandemic of SARS-CoV-2. Vaccination will remain the main measure for antiviral protection against COVID19 for a long time since the development of other types of antiviral drugs is much more time consuming [2].

At the same time, multiple recent studies have identified viral mutations that escape neutralizing antibodies targeting the SARS-CoV-2 Spike protein. Some of these mutations are already present in the human population [3] but many more may be present in natural reservoirs of coronaviruses and represent a potential threat [4,5]. These observations raise worries about the potency of monoclonal antibodies as well as the protective efficacy of the existing vaccines [6].

Great efforts have been undertaken by the scientific community in order to map potentially hazardous mutations [7]. Particularly, several sites at the SARS-CoV-2 Spike protein, which reduce the neutralizing activity of monoclonal antibodies and/or their cocktails/human sera were identified, including E484K, K417N [3], N439K [8], E406W [7], N501Y [9]. Many of these mutations occur in the receptor-binding domain (RBD) of Spike, which mediates binding to the angiotensin-converting enzyme 2 (ACE2) receptor resulting in the virus entry into the cells. At the same time, the majority of leading anti–SARS-CoV-2 antibodies also target this domain [10,11] rendering these mutations especially risky.

Due to the central role of RBD domain as a key Ab target, we have carried out a comprehensive computational investigation of the effects which may be induced by its mutations on the affinity to various neutralizing antibodies and hACE2 exploiting structural data available to date. We have assessed the impact of naturally occurring residue replacements in RBD to possible antibody resistance and the ACE2 binding as well as we analyzed the potential outcomes of all RBD mutations. We believe that the thorough virtual mutagenesis analysis reported here will guide further experimental studies of Ab resistance and identification of natural SARS-CoV-2 variants capable of antigenic escape.

## Materials and Methods

### Analysis of atomic structures and clustering

Structures of all RBD-Ab complexes were retrieved from the PDB database (see Table 1). In order to classify Ab epitopes, each Ab-RBD interface was encoded as a binary vector with the length equal to the number of residues in the reference RBD structure (in complex with the ACE2 receptor, the PDB code 6M17). Positions corresponding to residues in contact (distance between any pair of heavy atoms less than 6 Å) with an Ab were set to 1, while those not forming contacts were set to 0. The encoded epitopes were further clustered by means of the hierarchical algorithm and split into 4 clusters based on the inspection of the inter-cluster distances. The PDB structure 6M17 of the Spike protein in complex with the ACE2 receptor resolved by Cryo-EM to 2.90 Å [12] was used to estimate the effects of mutations on the RBD binding to ACE2.

**Table 1.**
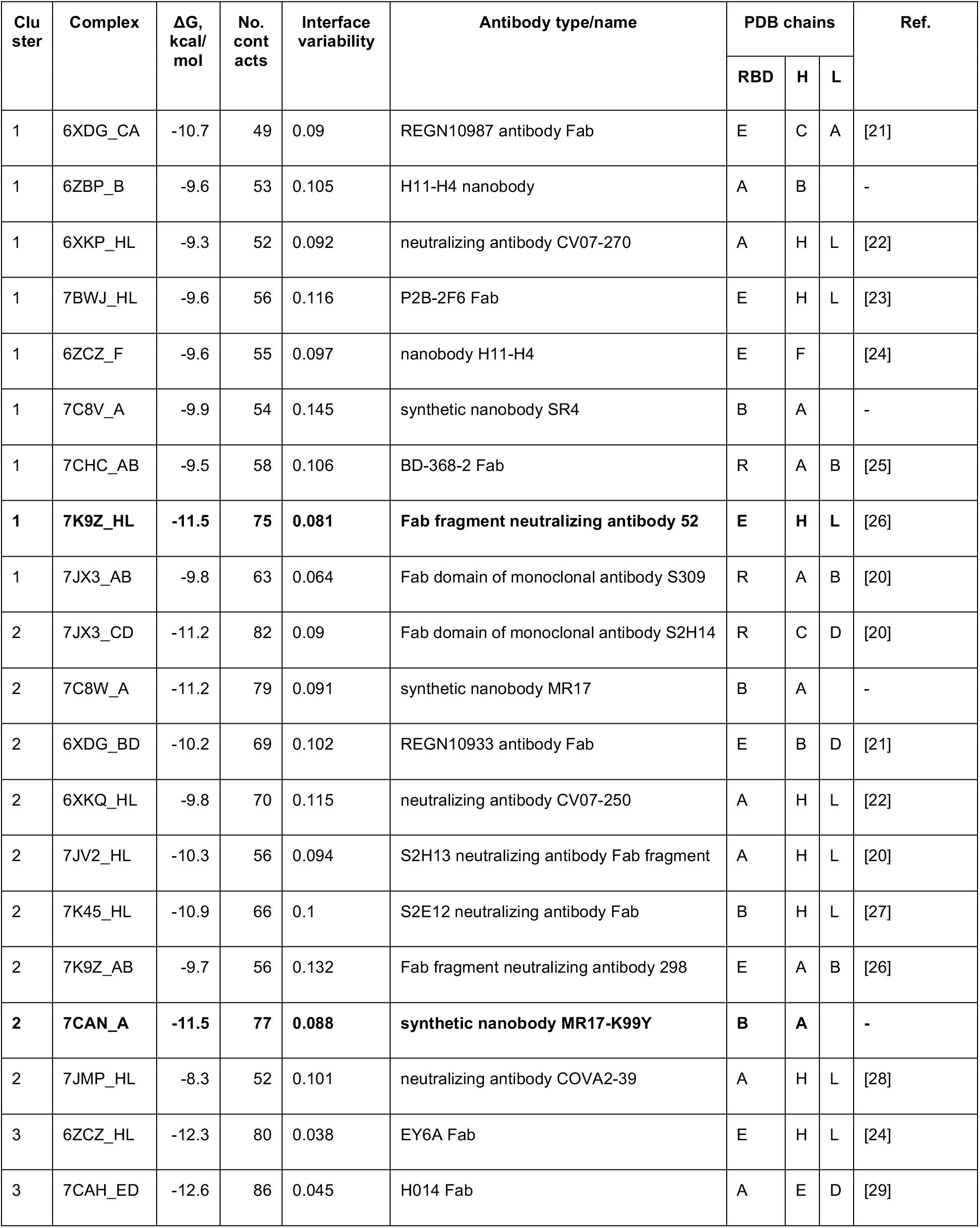
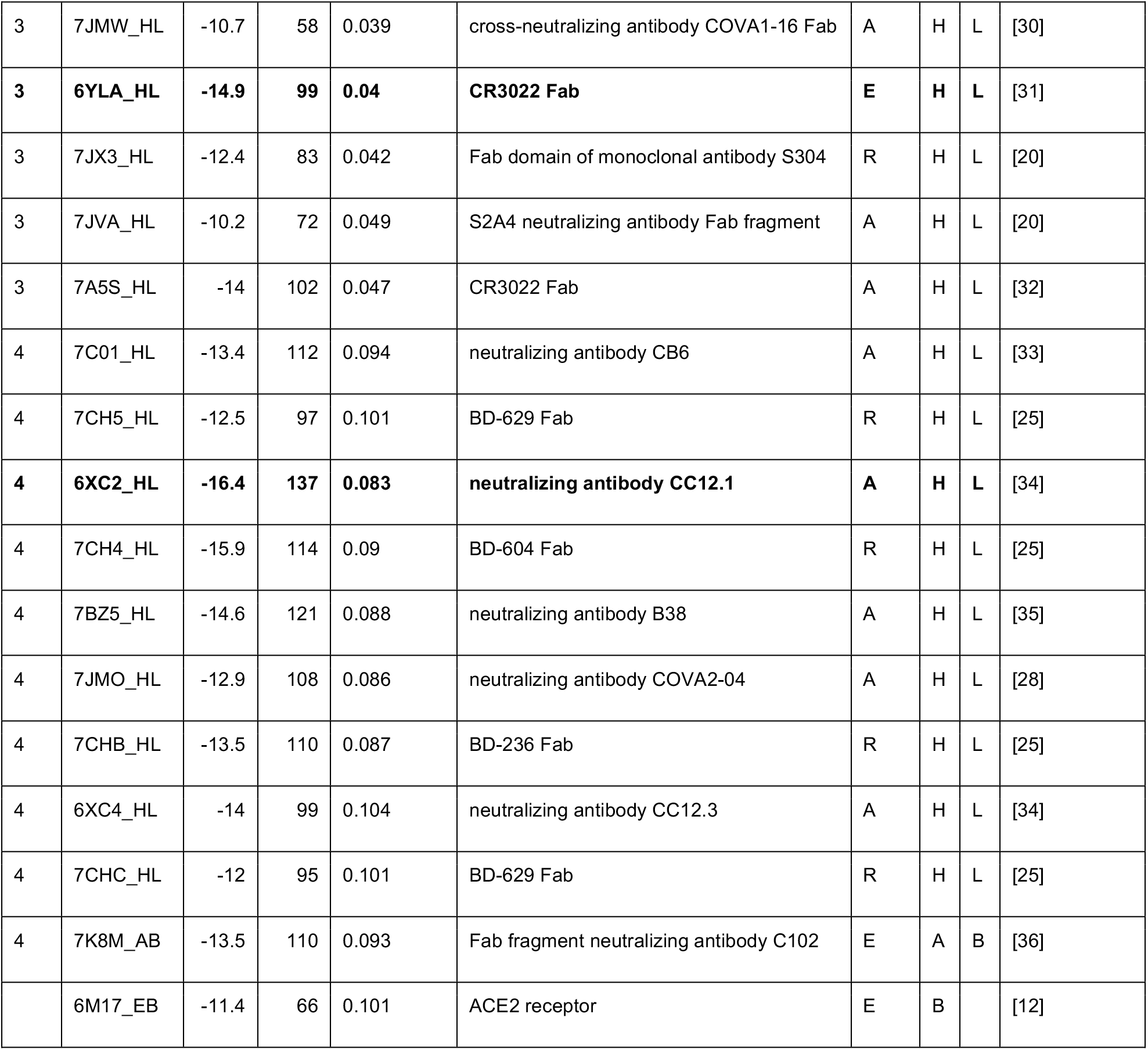
Overview of the analyzed antibody-RBD complexes. Complexes, which are centers of the clusters in Figure 1, are highlighted in bold. Some structures represent complexes with multiple Abs therefore for each complex the specific protein chains corresponding to RBD and Ab heavy/light chains are indicated (only one chain is shown for nanobodies).

### Estimation of binding energies

For calculation of binding energies between Abs, ACE2 and RBD, we used PRODIGY [13,14]. The contributions of amino acid mutations to the binding energies were estimated using the BeAtMuSiC server based on application of a statistical model to coarse-grained models of protein-protein complexes [15]. In order to estimate the effects of multiple mutations, the contributions made by individual ones were summed up.

### Sequence analysis

The sequences of the variants of the SARS-CoV-2 Spike protein were obtained from a database supported by the GISAID initiative [16] accessed on January/27/2021. A multiple sequence alignment (MSA) was obtained using MAFFT version 7 [17] with the default settings and truncated to the region corresponding to the RBD (C336-L518) present in the reference structure (PDB code 6M17) for further analysis. Specifically, the identical RBD sequences were sorted out and the variability of MSA positions was estimated in terms of Shannon entropy using ProDy [18]. In turn, the interface variability was estimated as the average entropy of amino acid positions forming the interface.

## Results and Discussion

### Analysis of epitopes reveals distinct clusters of Ab binding poses

The majority of human neutralizing antibodies (Ab) against SARS-CoV2 S-protein bind to diverse epitopes at the surface of its receptor-binding domain (RBD) preventing its attachment to the host cell and thus neutralizing the virus [19]. In order to explore the plasticity of their binding modes and classify them on the basis of residues involved into the complex formation with RBD, we applied the hierarchical clustering to the binding interfaces as described in the Methods. The analysis was performed for 35 Ab-RBD complexes available in the PDB database (Table 1, surveyed on December/14/2020). We further considered four major clusters of epitopes based on the distances observed in the clustering dendrogram (Fig. 1A). A representative complex for each cluster was chosen based on the estimated binding free energy. In each cluster, the Ab-RBD complex with the lowest binding free energy was picked up (hereafter referred by their PDB codes with the prefix standing for the Ab chains in the corresponding structure: 7K9Z_HL (cluster 1, cyan), 7CAN_A (cluster 2, yellow), 6YLA_HL (cluster 3, blue), and 6XC2_HL (cluster 4, red)). It may be noted also that complexes with lower binding energy Abs tend to form more contacts to RBD (see Fig. S1, Fig. S2) implying that focusing our analysis specifically on them should also allow us to examine the corresponding clusters of epitopes in a more complete way in addition to being stuck to potentially the most affine Abs.

**Figure 1.**
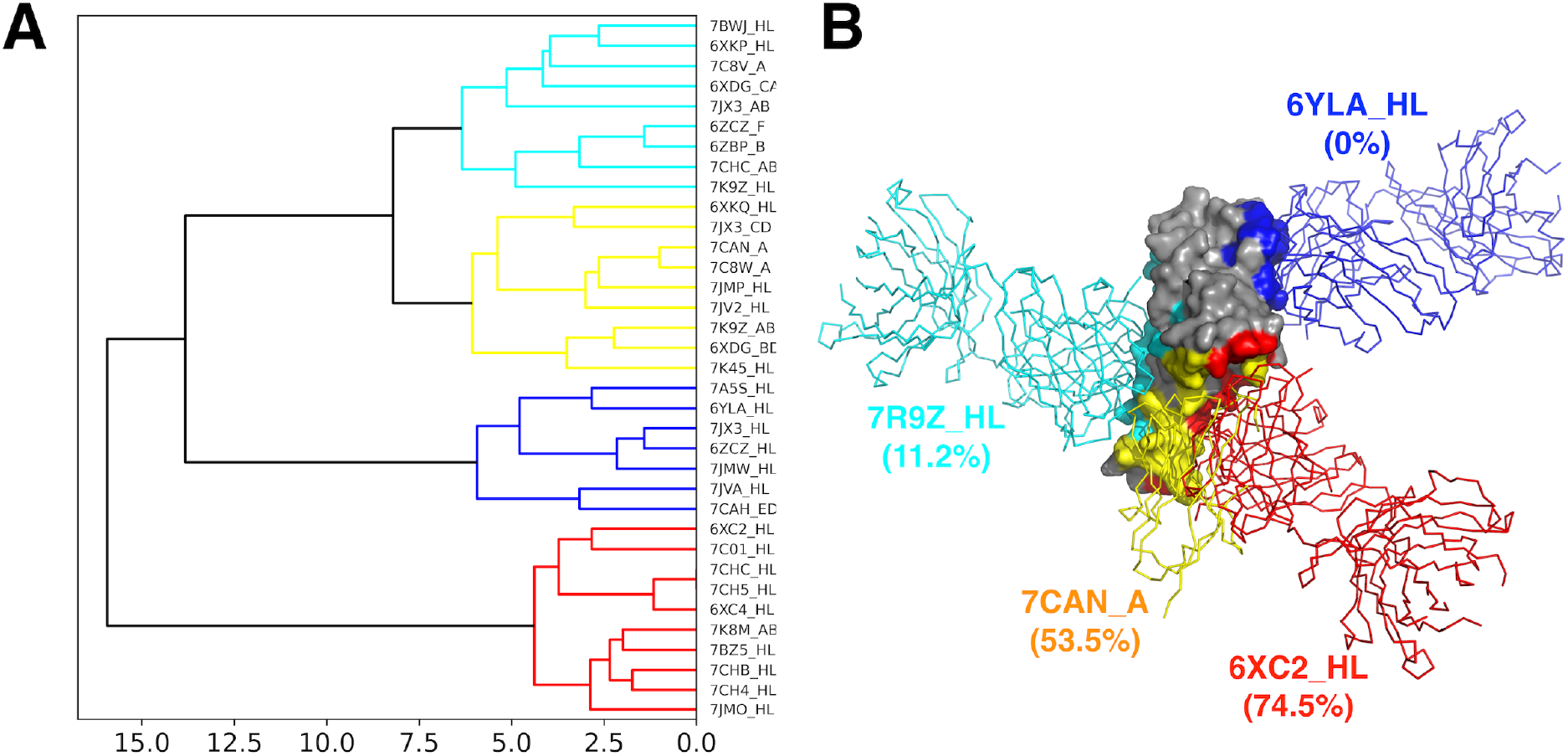
Cluster analysis of antibody epitopes. **A:** Hierarchical clustering dendrogram of binding epitopes. Four major clusters of epitopes are colored in cyan, yellow, blue, and red. **B:** Structures of representative antibody-RBD complexes for each cluster captioned with its PDB code. Colors correspond to panel A. Epitope overlaps with the ACE2 binding site are given in the parenthesis.

Detailed inspection of the interfaces in each cluster (Figure 1B) suggests that three of them extend over the same region of RBD encompassing the receptor-binding motif (RBM) β-hairpin of the RBD and partially overlap with each other and with the ACE2 binding site (clusters 1, 2, and 4; the average overlap with the ACE2 site is 11.2, 53.5, and 74.5%, respectively). However, the fourth cluster (cluster 3) occupies a distinct area on the RBD surface including residues 368-388 forming two α-helices and an intervening β-strand. This cryptic epitope is remote from the ACE2 site and not overlapping with it or epitopes of other clusters of Abs at all. While the Abs binding to this site do not prevent the RBD interaction with ACE2 in the direct way they sterically clash with ACE2 interacting with the same protomer within an S trimer [20].

Analysis of the amino acid variability of the SARS-CoV2 Spike RBD sequences available to date reveals that clusters 1, 2, and 4, which are spatially adjacent, exhibit significant level of variability while the epitopes of Abs belonging to cluster 3 are more conserved (Fig. 2, Fig. S2).

**Figure 2.**
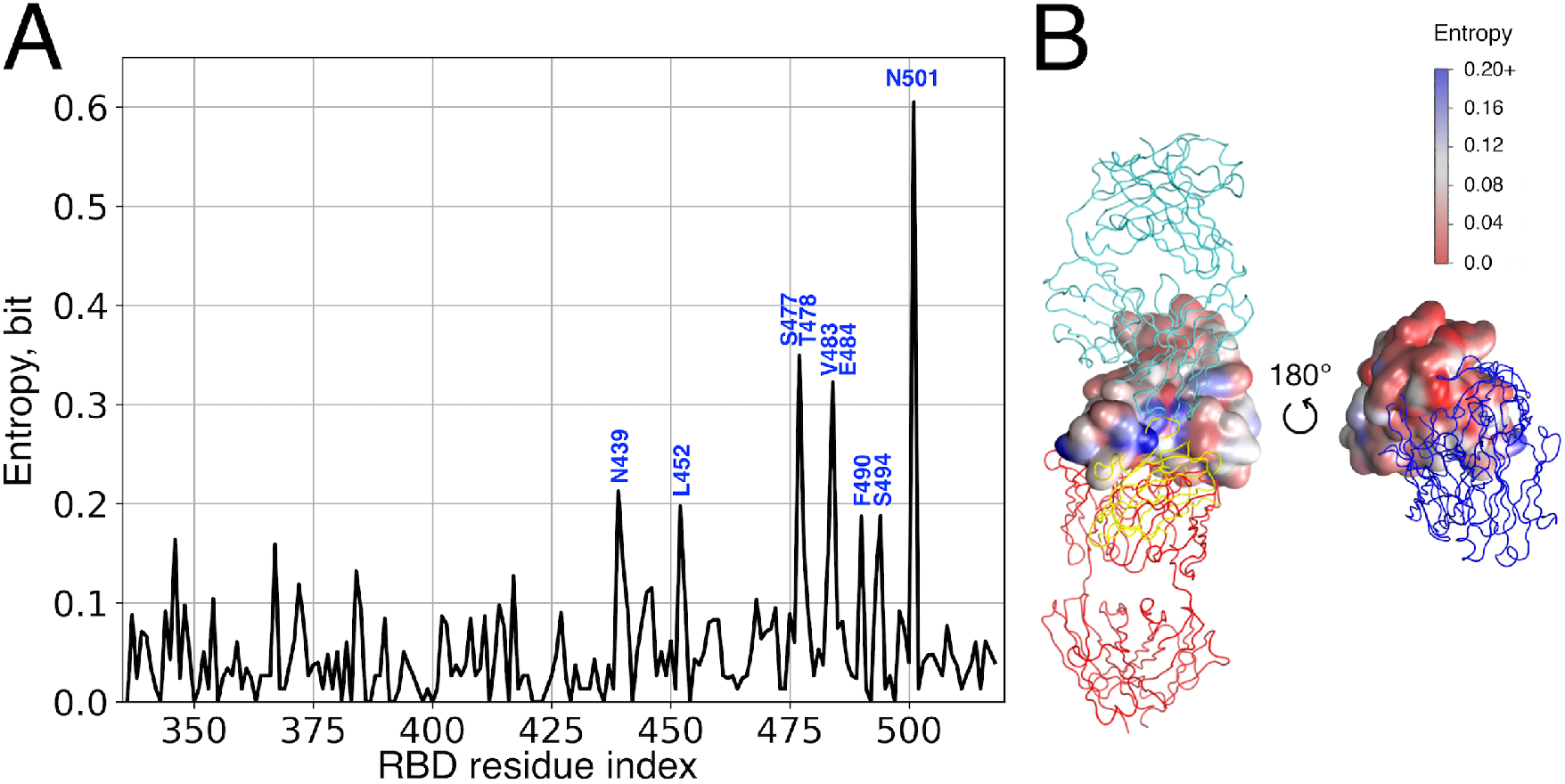
Sequence variability of RBD domain of S-protein. **A:** Shannon entropy (SE) calculated from the multiple sequence alignment (MSA) of RBD; residues with the high SE values are signed; **B:** Sequence variability in terms of SE mapped onto the surface of RBD. The positions of the four investigated antibodies are shown with respect to RBD.

### Effects of frequent RBD mutations on its binding affinity to Abs and ACE2

We further employed virtual mutagenesis analysis in order to assess the potential effects of point mutations in RBD on its binding affinity to antibodies belonging to four identified clusters and to the SARS-CoV-2 natural receptor, ACE2. Firstly, we estimated the effects on the binding energy for those single amino acid substitutions which are the most common mutations in the reported viral genomes. While almost all of them (Figure 3) appear to decrease the binding affinity to either all studied Abs (e.g., N439K) or some of them (e.g., E484K), this effect is relatively low. Notable exceptions are the G446V and K417N RBD variants, for which the binding energy is lowered by 1.3 kcal/mol and 0.9 kcal/mol towards 7CAN (cluster 2) and 6XC2 (cluster 4) Abs, respectively. At the same time, most mutations also result in lower affinity to ACE2. Remarkably, the most spread RBD mutation in the population to date, N501Y, which is particularly specific for B.1.1.7 lineage of variants, which are reportedly responsible for the recent outbreak in the UK, either does not cause any changes in affinity (to 7K9Z_HL, 7CAN_A, and 6YLA_HL Abs) or even results in a slight increase of the affinity (to 6XC2_HL Ab and the native receptor, ACE2). It is also worth to mention that affinity of 6YLA_HL, the representative Ab of cluster 3, seems to be the least affected by the analyzed mutations compared to Abs representing the other three clusters. This is apparently due to the remoteness of its epitope from the most frequent mutations, which mainly occur in the RBM region.

**Figure 3.**
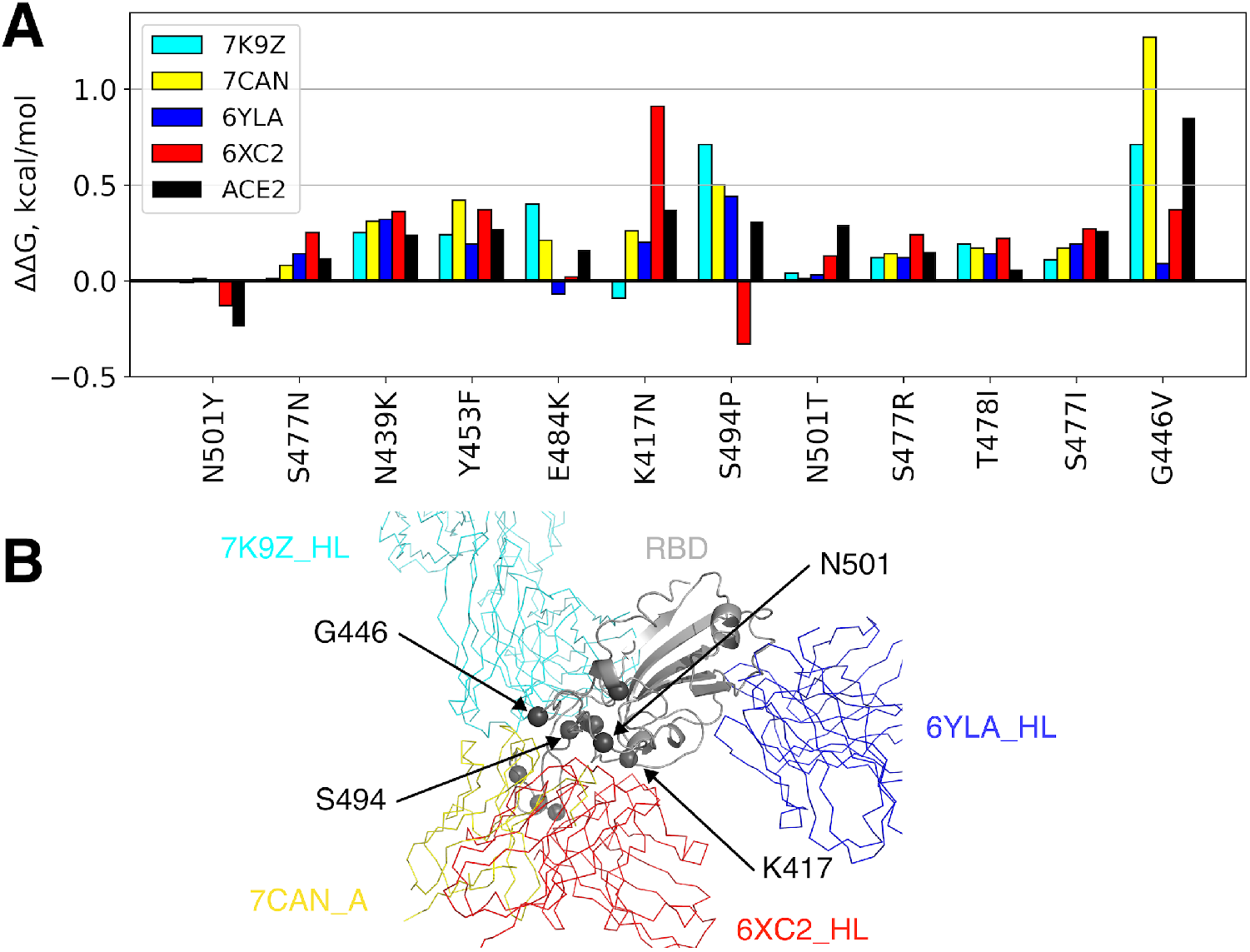
Influence of widely spread individual amino acid replacements on the affinity between antibodies/ACE2 and RBD. **A:** Predicted alteration of the binding free energy for the 12 most spread in the population RBD mutations; **B:** Locations of the mutated residues at RBD along with the spatial orientation of the 4 representative antibodies are shown.

Overall, these results suggest that none of the currently widely spread mutations of RBD are able to break the binding between RBD and the broad spectrum of Abs targeting it completely. However, two mutations may lead to partial loss of affinity to some of the analyzed Abs, such as K417N in case of 6XC2 and G446V in case of 7CAN. The former replacement, which is specific for the B.1.351 lineage (known as the South African variant) has been already shown to reduce the affinity to sera/monoclonal Ab [37].

### Computational mutagenesis predicts potentially deleterious RBD point mutations

Since CoVs undergo continuous and extensive antigenic evolution [38], which can potentially lead to numerous amino acid variations that are not currently observed but can appear in future, we did not limit our analysis to the existing variants but performed the complete virtual mutation scanning of RBD (notably, the least conserved domain of Spike [8] by systematically changing all amino acid positions to 19 alternatives. The latter task was allowed here by exploiting the fast computational approach for the virtual mutagenesis.

The effects of the majority of mutations on the binding affinity is very low as indicated by corresponding distributions of the predicted ΔΔG values (Fig. 4, middle panel). However, all of these distributions are not symmetrical but have a longer right tail corresponding to positive ΔΔG values and, thus, the decrease of affinity. In other words, the number of mutations that lower affinity and the extent of this lowering effect is greater than for mutations that strengthen the binding. This observation is also true for the RBD-ACE2 interaction (see Fig. 4N).

**Figure 4.**
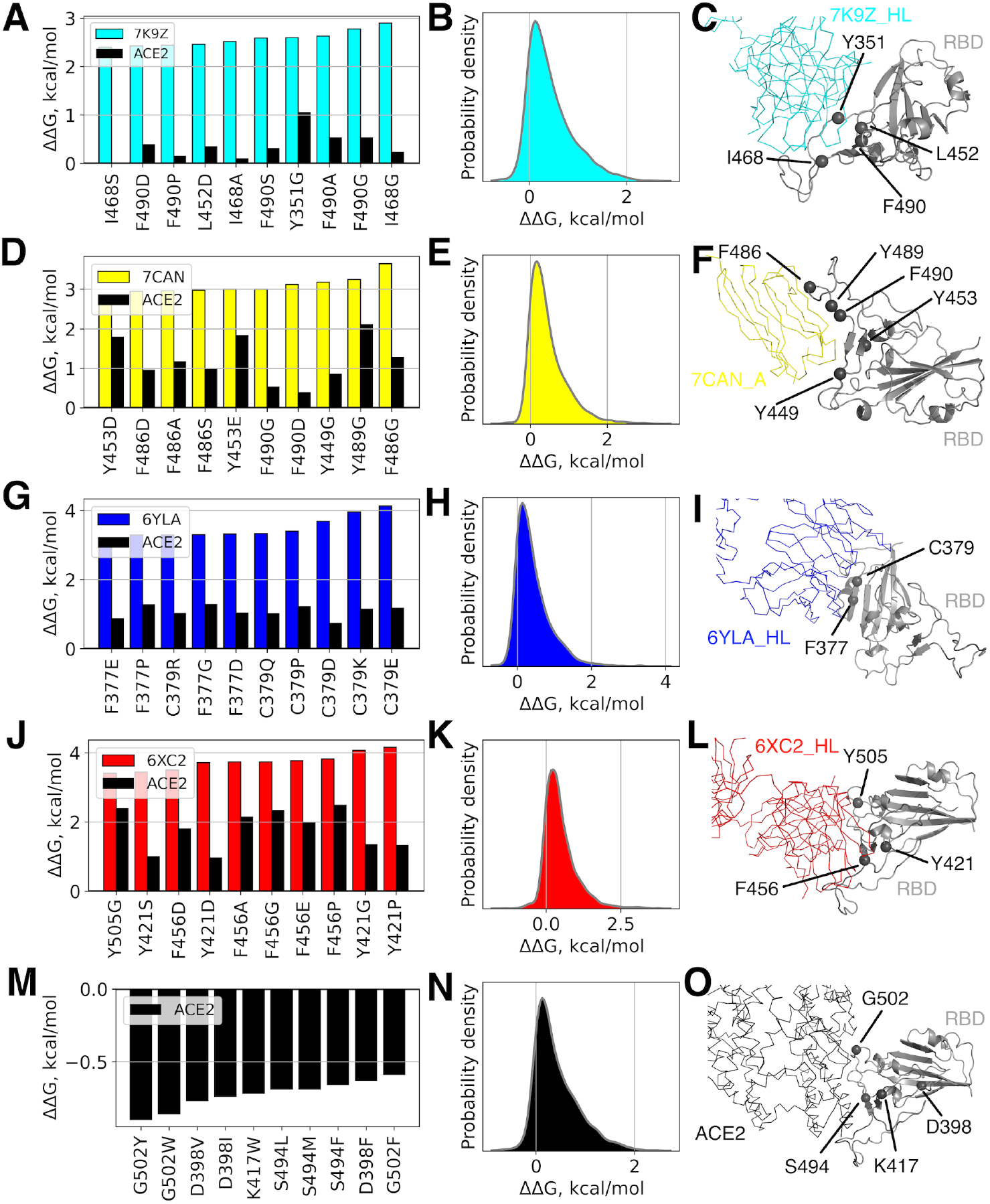
Effects of single mutations on the binding affinity of the RBD-antibody and RBD-ACE2 complexes. RBD mutations with the largest positive (for different antibodies, **A, D, G, J**) and negative (for ACE2, **M**) predicted contributions are shown in the left panels; positions of these amino acids in structures of the corresponding protein complexes are shown in the right panels; the distributions of predicted affinity alterations caused by all the possible single mutations are shown in the middle panels.

Despite the very low effects induced by the vast majority of mutations on the RBD-Ab/ACE2 binding energy, some mutations may potentially change it by up to +3-4 kcal/mol, what might already reduce the affinity between proteins drastically given the typical Ab-RBD binding energies (see Table 1, ΔG_average_=−11.7 kcal/mol; experimental values fall into the same range, e.g. for CR3022 K_D_=~6.6 nM - ~115 nM [31,39,40] corresponding to ΔG=−11.6 - −9.8 kcal/mol; for ACE2 - KD=~15nM [10], i.e. ΔG=-11.1 kcal/mol at 310 K).

Importantly, mutations with strong effects on the binding energy usually occur directly or in a close vicinity of an Ab epitope often resulting in the reduced affinity of RBD to ACE2 as well as for the majority of Abs their interfaces overlap with the one of ACE2. We noticed certain correlation trend between ΔΔGs estimated for mutations in the RBD-ACE2 complex and ΔΔGs for the same mutations in the all of studied Ab-RBD complexes (see Fig. 5, Pearson’s r=0.59÷0.81). However, it is more pronounced for the 6XC2 Ab, whose interface overlaps with ACE2 the most (~75%), and less for other Abs. Still, a number of mutations reduce the RBD affinity towards Abs without any noticeable change (or even increased affinity) towards ACE2. Such mutations are listed in Table S1.

**Figure 5.**
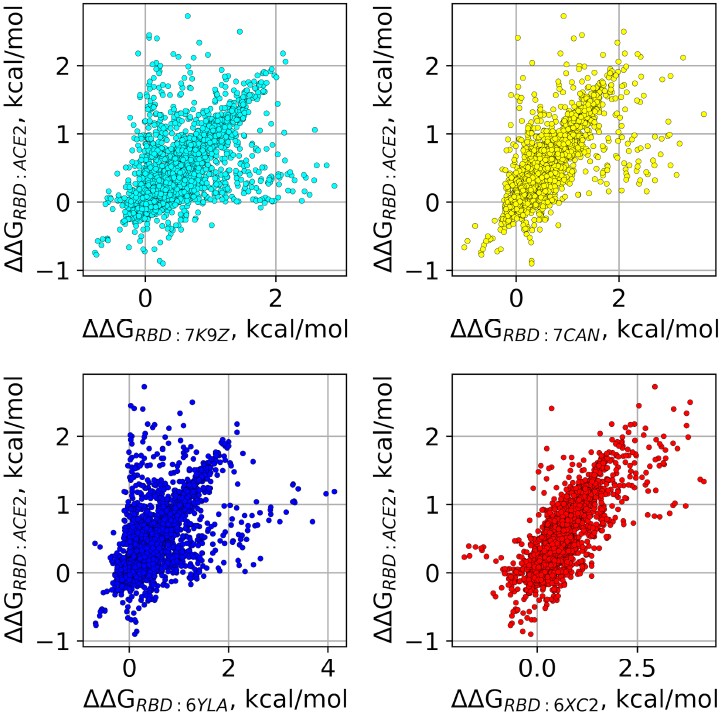
Analysis of single amino acid RBD mutants. Affinity changes of single amino acid RBD mutants to different antibodies are plotted in comparison to the respective affinity changes to ACE2.

It is important to say that some of crucial mutations may also seriously destabilize the structure of RBD or even that of the whole S-protein rendering them non-functional/incompetent of the ACE2 binding [7]. While the complete evaluation of such destabilizing effects is of greater complexity and is beyond the scope of the present work, we can still draw some conclusions without providing an additional analysis. For instance, the RBD mutations which affect the 6YLA binding the most (see Fig. 4G) involve C379 forming a disulfide bridge to C432. Thus, all of these mutations are likely to influence the RBD structure and reduce its affinity to Ab. Nonetheless, the major part of the impactful mutations is not expected to impair the overall RBD stability and thus may be considered of a high risk for a potential antigenic escape.

### Antigenic escape assessment for existing SARS-CoV2 variants

Finally, we estimated potential changes of the binding energy between RBD and four studied Abs caused by multiple mutations co-occurring in the real viral sequences stored at GISAID to date. The total affinity change was estimated as a sum of the binding energy changes induced by all the individual mutations in a given sequence and compared with the cognate change of the RBD affinity towards ACE2.

In general, the results indicate that higher number of mutations in RBD quite expectedly leads to larger ΔΔG (see Fig. 6A, Pearson’s r=0.57÷0.84). At the same time, as few as 2 substitutions are sufficient to reduce RBD affinity to individual Abs by up to 2.5-3.0 kcal/mol, while 6 substitutions may result in ΔΔG up to 3.3-4.0 kcal/mol. The SARS-CoV2 variants with the largest estimated change in RBD-Ab/ACE2 affinity are listed in Table 2. Among them, the most drastic decrease of affinity was predicted for 7CAN and 6XC2 Abs towards the RBD variants found in animals, bats and pangolins. These RBD sequences comprise as many as 20-25 mutations. At the same time, the largest decrease in the affinity towards 7K9Z and 6YLA Abs was estimated for the two sequences obtained from human samples in Iran and accommodated 8 and 11 substitutions, respectively. It is also worth mentioning that all of the variants with the largest affinity decrease towards particular Ab also demonstrate decreased to some extent (at least by ~2.1 kcal/mol) affinity towards other examined Abs. The same is true for their affinity to ACE2 which is dropped by 2.7-6.9 kcal/mol. Furthermore, a correlation trend between ΔΔG for RBD-Ab and ΔΔG for RBD-ACE2 can be noted (Fig. 6B, Pearson’s *r*=0.63-0.90) similar to the correlation observed for single mutations and discussed in the previous section. It does not exclude, however, the presence of some RBD variants for which the affinity towards specific Abs is decreased by ~2-4 kcal/mol while there is no notable change of affinity to ACE2 (Fig. 6B), although for 6XC2 Ab the effect is lower and does not exceed 2 kcal/mol. The latter fact is apparently due to the largest overlap of the 6XC2 Ab epitope with the ACE2 interface making those RBD mutations, which are unfavorable for the binding of this particular Ab, also adverse for the RBD interaction with ACE2. On the other hand, one cannot exclude a possibility that variants escaping the 6XC2 Ab and, at the same time, interacting with ACE2 in the normal way can appear in future. The analysis of single mutations performed above indicates (see Table S1) that as many as six point mutations may potentially result in the substantial decrease of the RBD affinity to the 6XC2 Ab by ~5.6 kcal/mol while the affinity of this variant to ACE2 would even increase by ~0.4 kcal/mol.

**Figure 6.**
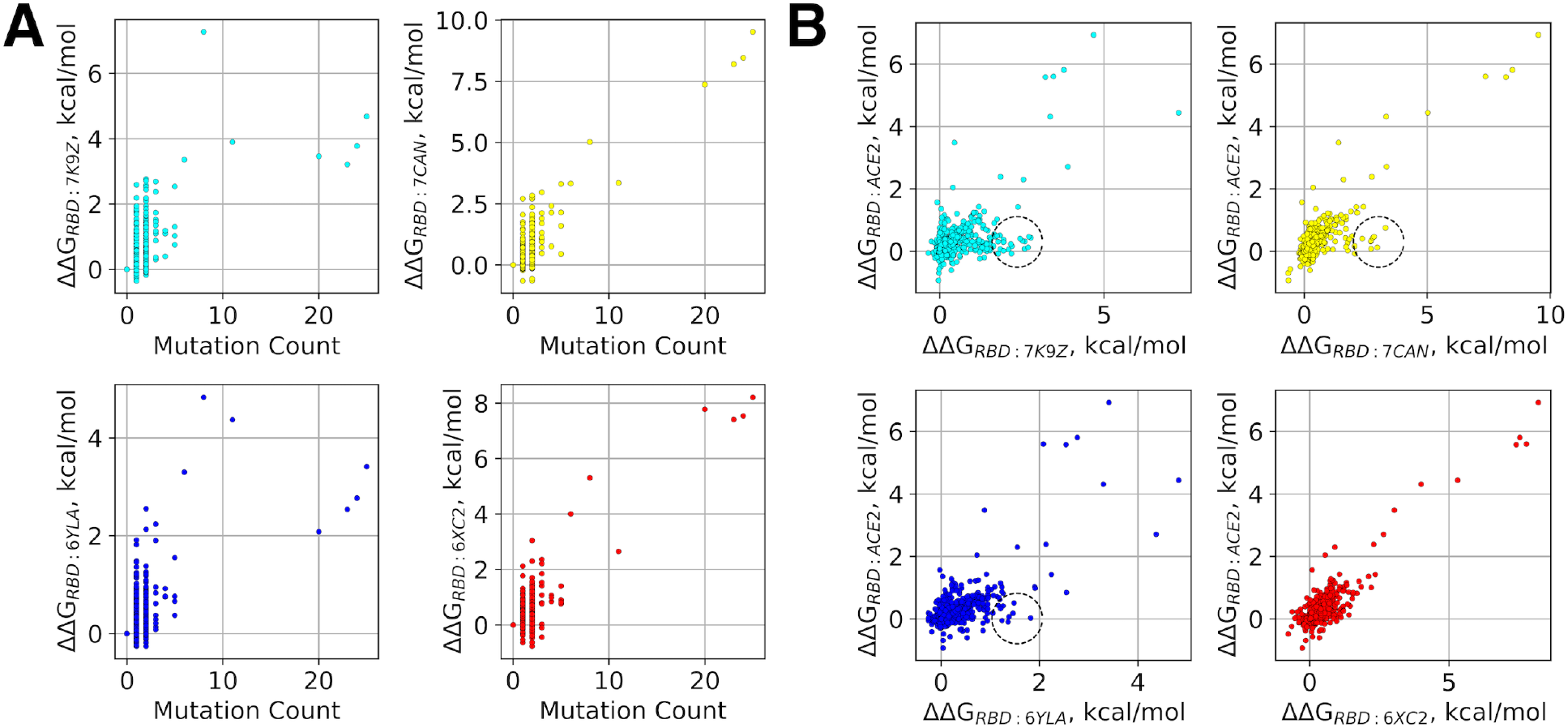
Analysis of RBD variants. **A:** Affinity changes of RBD variants to different antibodies as a function of the total count of mutations in an RBD mutant; **B:** Affinity changes of RBD variants to ACE2 are plotted as a function of the respective affinity changes to different antibodies. Dashed cycles indicate RBD variants with significant decrease of affinity towards antibodies, which is not accompanied by any notable change of affinity to ACE2.

**Table 2.** Existing SARS-CoV2 variants with the largest predicted change of the RBD-Ab/ACE2 binding energy. Variants with the largest positive change are provided for different antibodies (ΔΔG > 0 resulting in lower affinity to Abs), while for ACE2 the variant with the largest negative change is listed (ΔΔG < 0 resulting in higher affinity to ACE2).

| ID | Origin | No. mutations | $\Delta\Delta G$, kcal/mol | | | | |
| --- | --- | --- | --- | --- | --- | --- | --- |
|  |  |  | RBD:7K9Z | RBD:7CAN | RBD:6YLA | RBD:6XC2 | RBD:ACE2 |
| EPI_ISL_410541 | pangolin/Guangxi | 25 | 4.68 | **9.52** | 3.41 | 8.21 | 6.93 |
| EPI_ISL_410538 | pangolin/Guangxi | 24 | 3.78 | **8.46** | 2.77 | 7.54 | 5.81 |
| EPI_ISL_410539 | pangolin/Guangxi | 23 | 3.21 | 8.2 | 2.54 | **7.41** | 5.58 |
| EPI_ISL_402131 | bat/Yunnan | 20 | 3.46 | 7.37 | 2.08 | **7.78** | 5.6 |
| EPI_ISL_568499 | human/Iran | 8 | **7.27** | 5.02 | 4.83 | 5.31 | 4.44 |
| EPI_ISL_568500 | human/Iran | 11 | 3.9 | 3.35 | **4.37** | 2.65 | 2.71 |
| EPI_ISL_802528 | human/Netherlands | 2 | -0.04 | -0.64 | -0.64 | -0.27 | **-0.93** |

Finally, it is noteworthy to point to a SARS-CoV2 variant with the largest increase of affinity towards ACE2 (~0.9 kcal/mol, Table 2), which was obtained from a human sample and bears just 2 substitutions in the RBD region. This affinity change co-occurs together with a slight increase of an affinity towards all examined Abs ranging from almost zero to 0.6 kcal/mol.

## Conclusion

The enormous rates, at which the scientific community is studying SARS-CoV-2, have generated a huge amount of data about the novel coronavirus since its emergence about a year ago. The open databases of sequences and atomic structures provide an opportunity to perform the high-throughput analysis of interactions between various monoclonal neutralizing antibodies (Abs), hACE2, serving as the main entry point for coronavirus, and their principal viral binding partner, the receptor-binding domain (RBD) of Spike. Here, we employed a theoretical approach to predict the effects of the known and conceivable RBD mutations on the binding energy between RBD, ACE2 and a set of representative Abs targeting structurally distinct epitopes.

Analysis of epitopes revealed four major clusters of Ab binding sites differing in both involved RBD residues and their conservativity. While most of the currently widely spread mutations appear in the least conservative part of RBD, receptor-binding motif (RBM), we show that there also exist potentially deleterious mutations in more conservative regions of RBD, which can significantly impair its binding to Abs. While certain point replacements in RBD, including those, which has been previously characterized experimentally and/or observed in the population, can seriously reduce binding affinity for specific Abs or even several types of Abs, the synergic effects of several multiple mutations may be much more hazardous leading to resistance against a broad spectrum of Abs. By analyzing sequences of the existing SARS-CoV-2 variants, we have identified a number of such naturally occurring mutants potentially capable of the extensive antigenic escape. Moreover, we have identified several mutations which, unlike the majority of replacements, are predicted to reduce binding affinity between RBD and one or several Abs apparently without weakening the RBD-ACE2 binding. Such mutations may be especially threatening in terms of the emergence of resistant and highly contagious variants of SARS-CoV-2.

We expect that our theoretical study will promote future experimental work and complement it allowing eventually to prognose properties of novel SARS-CoV-2 variants and to design more efficient treatments for COVID-19. Particularly, the knowledge-based selection of antibody mixtures with non-overlapping escape mutations should reduce the emergence of resistance and prolong the utility of antibody therapies.

## Acknowledgments

The study has been supported by the RFBR grant #20-04-60258 to O.S.S.

M.P.K., O.S.S., K.V.S. acknowledge the support from the Interdisciplinary Scientific and Educational School of M.V. Lomonosov Moscow State University «Molecular Technologies of the Living Systems and Synthetic Biology».

