## Supplementary Materials for "Effects of Mutations in the Receptor-Binding Domain of SARS-CoV-2 Spike on its Binding Affinity to ACE2 and Neutralizing Antibodies Revealed by Computational Analysis"

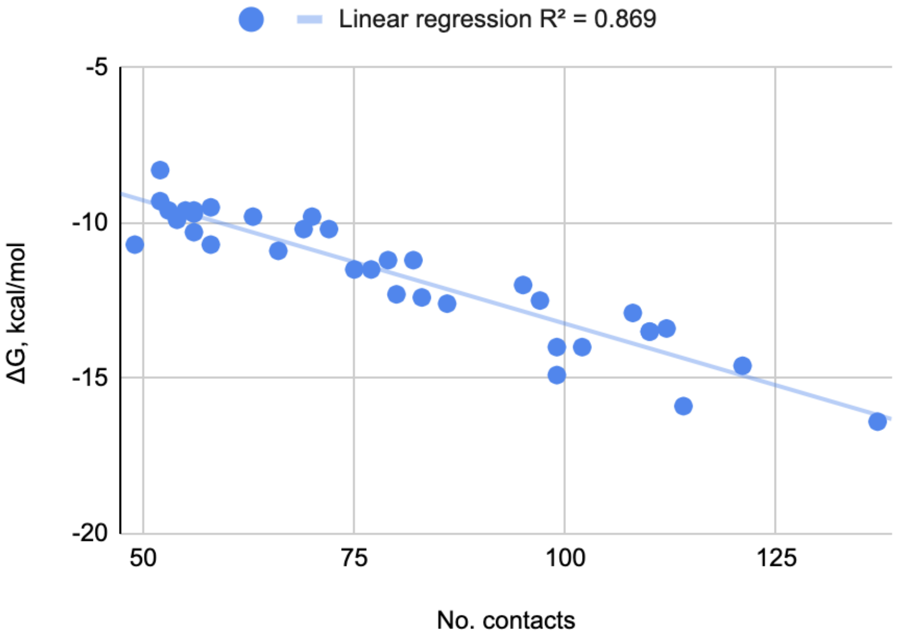


**Figure S1.** Theoretical estimation of the free energy of the Ab-RBD complexes plotted as a function of the contact count. The linear regression trend is shown.

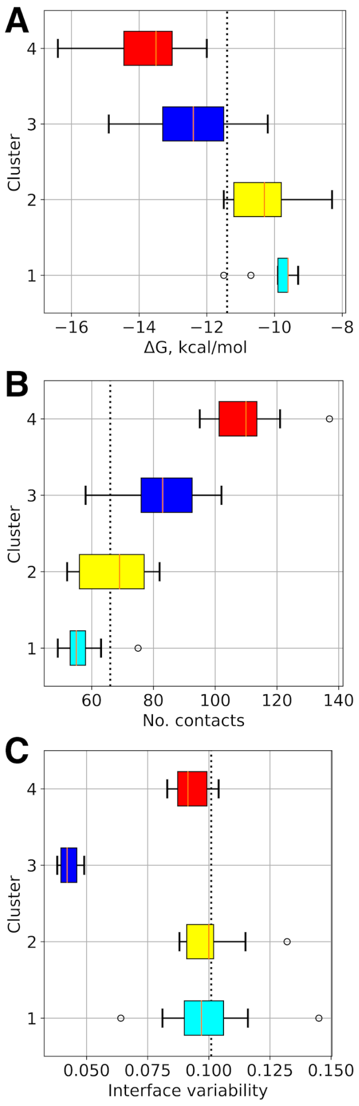

**Figure S2.** Box plots of the binding free energy (A), number of contacts between Ab and RBD (B); and interface variability in terms of average entropy (C) for each cluster of analyzed Ab-RBD complexes.

**Table S1.** List of mutations, which lower the binding affinities between RBD and antibodies (ΔΔG > 0.6 kcal/mol) but do not lower the binding affinity to ACE2 (ΔΔG ≤ 0 kcal/mol).

| **Antibody** | **Variant** | **ΔΔG, kcal/mol (RBD:Ab)** | **ΔΔG, kcal/mol (RBD:ACE2)** |
|---|---|---|---|
| 7K9Z_HL | P337M | 0.62 | -0.22 |
| 7K9Z_HL | P337R | 0.62 | -0.26 |
| 7K9Z_HL | P337E | 0.63 | -0.16 |
| 7K9Z_HL | P337Q | 0.63 | -0.19 |
| 7K9Z_HL | P337G | 0.64 | -0.06 |
| 7K9Z_HL | V483C | 0.66 | -0.01 |
| 7K9Z_HL | P337Y | 0.76 | -0.26 |
| 7K9Z_HL | P337F | 0.77 | -0.24 |
| 7K9Z_HL | P337W | 0.81 | -0.15 |
| 7K9Z_HL | I468F | 0.81 | 0 |
| 7K9Z_HL | I468H | 0.88 | -0.02 |
| 7K9Z_HL | G482C | 0.89 | 0 |
| 7K9Z_HL | V483G | 1.02 | -0.34 |
| 7K9Z_HL | I468T | 1.77 | -0.08 |
| 7K9Z_HL | I468N | 1.83 | -0.01 |
| 7CAN_A | E484S | 0.61 | 0 |
| 7CAN_A | S383P | 0.65 | -0.05 |
| 7CAN_A | P337W | 0.67 | -0.15 |
| 7CAN_A | V483G | 0.82 | -0.34 |
| 7CAN_A | E484A | 0.83 | -0.01 |
| 6YLA_HL | V362A | 0.64 | 0 |
| 6YLA_HL | D428T | 0.66 | -0.04 |
| 6YLA_HL | P337K | 0.68 | -0.18 |
| 6YLA_HL | P337E | 0.7 | -0.16 |
| 6YLA_HL | P337G | 0.72 | -0.06 |
| 6YLA_HL | Y369L | 0.72 | -0.1 |

| | | | |
|---|---|---|---|
| 6YLA_HL | K378T | 0.76 | -0.19 |
| 6YLA_HL | T385N | 0.85 | -0.13 |
| 6YLA_HL | K378Q | 0.88 | 0 |
| 6YLA_HL | K378C | 0.93 | -0.12 |
| 6YLA_HL | D428A | 0.95 | 0 |
| 6YLA_HL | K378S | 1.21 | -0.07 |
| 6YLA_HL | T385E | 1.23 | -0.09 |
| 6YLA_HL | K378N | 1.24 | -0.05 |
| 6YLA_HL | T385D | 1.24 | -0.17 |
| 6YLA_HL | T385Q | 1.24 | -0.02 |
| 6YLA_HL | G381A | 1.25 | -0.04 |
| 6YLA_HL | G381S | 1.26 | -0.07 |
| 6YLA_HL | Y369I | 1.33 | -0.02 |
| 6YLA_HL | S383P | 1.35 | -0.05 |
| 6YLA_HL | T385S | 1.36 | -0.09 |
| 6YLA_HL | T385P | 1.5 | -0.05 |
| 6XC2_HL | N481P | 0.64 | -0.11 |
| 6XC2_HL | R403L | 0.66 | -0.04 |
| 6XC2_HL | P337K | 0.67 | -0.18 |
| 6XC2_HL | R403I | 0.67 | -0.01 |
| 6XC2_HL | P337E | 0.68 | -0.16 |
| 6XC2_HL | N422P | 0.68 | -0.18 |
| 6XC2_HL | T415R | 0.75 | -0.03 |
| 6XC2_HL | P337G | 0.84 | -0.06 |
| 6XC2_HL | T415C | 1.01 | -0.02 |
| 6XC2_HL | G502R | 1.07 | -0.01 |
| 6XC2_HL | R403P | 1.36 | -0.05 |
